# SavvyCNV: genome-wide CNV calling from off-target reads

**DOI:** 10.1101/617605

**Authors:** Thomas W Laver, Elisa De Franco, Matthew B Johnson, Kashyap Patel, Sian Ellard, Michael N Weedon, Sarah E Flanagan, Matthew N Wakeling

## Abstract

Identifying copy number variants (CNVS) can provide diagnoses to patients and provide important biological insights into human health and disease. Current exome and targeted sequencing approaches cannot detect clinically and biologically-relevant CNVs outside their target area. We present SavvyCNV, a tool which uses off-target read data to call CNVs genome-wide. Up to 70% of sequencing reads from exome and targeted sequencing fall outside the targeted regions - SavvyCNV exploits this ‘free data’.

We benchmarked SavvyCNV using truth sets generated from genome sequencing data and Multiplex Ligation-dependent Probe Amplification assays. SavvyCNV called CNVs with high precision and recall, outperforming five state-of-the-art CNV callers at calling CNVs genome-wide using off-target or on-target reads from targeted panel and exome sequencing. Furthermore SavvyCNV was able to call previously undetected clinically-relevant CNVs from targeted panel data highlighting the utility of this tool within the diagnostic setting. SavvyCNV is freely available.

---

Copy number variants (CNVs) are an important class of biological variation. They can cause monogenic disease (1,2), are associated with polygenic traits (3) and may exert pharmacogenetic effects(4). CNVs are structural rearrangements where bases are gained (duplication) or lost (deletion) from the genome causing an altered copy number compared to the reference.

The importance of CNVs is highlighted by the role they play in many diseases, including cancers (5), autism (6), developmental disorders (7), and heart disease (8). Single or partial gene deletions can cause disease where haploinsufficiency would result in the disease phenotype. For example, both single nucleotide variants and whole gene deletions of *PKD1* cause polycystic kidney disease (1). Duplications can also cause disease as a result of gene disruption at the site of insertion or through increased gene expression. For example, paternal duplication of the chromosome 6q24 region causes neonatal diabetes by overexpression of the imprinted gene *PLAGL1* (2,9). Larger CNVs are likely to cause syndromic disease as they affect multiple genes. An extreme case is Down syndrome where duplication of chromosome 21 results in characteristic facial features and intellectual disability (10).

CNVs can be detected by a range of methods. In the clinical setting DNA microarrays are routinely used to detect larger rearrangements whilst multiplex ligation-dependent probe amplification (MLPA) is often used to detect single or partial gene CNVs (11). With next generation sequencing (NGS) increasingly employed to investigate genetic variation, the detection of CNVs from NGS data has become increasingly important. While genome sequencing is the optimal method to capture all sequence variation across the genome, due to speed and cost exome sequencing and targeted NGS panels are the most commonly used testing methods, particularly as a first line test in clinical diagnostic laboratories.

Many methods have been published to call CNVs from exome and targeted gene panel data(12). These are designed to detect CNVs within the genes which are targeted by the assay, however biologically interesting and disease causing CNVs will often fall outside of the targeted regions. Existing methods will typically be able to identify if a particular gene is deleted/duplicated however they will not necessarily be able to map the extent of the CNV as the breakpoints will often be located outside the targeted regions.

Current approaches to gene targeting for NGS are imperfect. Samuels *et al* reported that between 40% and 60% of sequence reads generated map outside the target regions (13). This in effect produces ultra-low depth genome sequence data. While there is insufficient information (<1X coverage) to call single nucleotide variants over the untargeted region this ‘off target’ data can be exploited to call large CNVs by detecting read depth changes over a wide area. The ability to use offtarget reads to call CNVs across the genome increases the diagnostic utility of targeted next-generation sequencing panels and also allows for more accurate mapping of breakpoints of CNVs which reside outside of the targeted regions. Previous tools have been designed to detect CNVs in off-target reads from exome data (14) and large targeted panels (15,16).

We have developed a new tool, SavvyCNV, for calling CNVs from off-target reads. We investigated the utility of SavvyCNV by comparing it to the current tools for calling CNVs in off-target regions in both targeted sequencing and whole exome sequencing (using a truth set derived from genome sequencing), and in on-target regions (using a truth set derived from MPLA). We then used SavvyCNV in a patient cohort tested with a small targeted gene panel (75 genes) to perform a genome-wide analysis to detect CNVs of clinical relevance.

## Material & Methods

### Targeted panel data

2591 patients were referred to the molecular genetics department at the Royal Devon and Exeter Hospital for genetic testing for maturity onset diabetes of the young (MODY), neonatal diabetes (NDM) or hyperinsulinemic hypoglycemia (HH). Informed consent was obtained from the probands or their parents/guardians and the study was approved by the North Wales ethics committee. The study was conducted in accordance with the Declaration of Helsinki. Samples were sequenced on a targeted gene panel test for monogenic diabetes and HH using a custom Agilent SureSelect panel of 75 genes, targeting 200kb and obtaining 3.4 million reads on average per sample (standard deviation 1.6 million)(17), using an Illumina HiSeq 2500 or an Illumina NextSeq 500. Based on the GATK best practice guidelines(18) reads were aligned to the hg19/GRCh37 human reference genome with BWA mem(19), duplicates were removed using Picard (https://broadinstitute.github.io/picard/) and GATK IndelRealigner was used for local re-alignment. CNV analysis was carried out on these BAM files.

### Exome sequencing data

Following testing using the targeted panel, samples from 86 patients underwent exome sequencing with Agilent SureSelect Whole Exome versions 1, 3, 4, and 5, obtaining 76 million reads on average (standard deviation 20 million). Sequencing, alignment and variant calling was as above for the targeted panel.

### Truth set for targeted and exome data

170 of the targeted panel samples and 42 of the exome samples were subsequently genome sequenced on an Illumina HiSeq 2500 or an Illumina HiSeq X10. These were used to create a truth set of CNVs for testing the off-target CNV calling from targeted panel or exome data. The CNVs in the truth set were called by GenomeStrip(20) from the genome sequencing data. In order to remove false positive calls CNVs were filtered based on their allele balance ratios – whether the allele balance of the variants within the called CNV was consistent with it being a true call. We used the X chromosome in males to calibrate the expected allele ratio for a deletion and used the allele ratio of normal, two copy regions to evaluate if the allele ratio for duplications fell above that. In addition 37 CNVs were added to the targeted panel truth set as they were validated by other methods such as Multiplex Ligation-dependent Probe Amplification (MLPA) or were aneuploidies reported by the clinician at time of referral for genetic testing.

The ICR96 data set (21) was used to benchmark on-target CNV calling. This data set consists of 96 samples sequenced on a targeted panel where the truth set of CNVs is based on 68 positive and 1752 negative MLPA tests.

### Calling clinically-relevant CNVs

The remaining 2479 targeted panel samples from unsolved patients with MODY, NDM and HH were analysed with SavvyCNV to look for off-target CNVs which might explain their phenotype. For clinical evidence of the CNVs, see Table 4.

### CNV tool comparisons

To ensure a fair comparison between the different tools, for each data set all tools were run with a variety of configurations. The size of genomic regions that were analysed was varied for all six tools (targeted panel: 150kbp to 300kbp or 50kbp to 2Mbp for CopywriteR; exomes: 6kbp to 50kbp or 20kbp to 2Mbp for CopywriteR; ICR96: 200bp to 600bp). The hidden Markov model transition probability was varied for DeCON and SavvyCNV (10^-10^ to 0.1). All six tools provide quality metrics for the CNV calls. These metrics were used to filter the CNV calls to reject false positive calls and retain true positive calls. All possible quality cut-off values were tried. The best precision achieved for each possible recall was then selected for each tool from all the generated results, and plotted in precision-recall graphs. EXCAVATOR2(14) did not run on the ICR96 data set - we contacted the authors of the tool but did not receive a response. CopywriteR was not run on the ICR96 data set, as it is not designed to use on-target data.

### Statistics

We defined recall as the percentage of true positive CNVs that were found by the tool. We defined precision as the percentage of the total CNVs called by the tool that were true. Several figures use the f statistic to compare tools; this is the harmonic mean of precision and recall.

## Results

### How much off-target read data is there?

For our small targeted panel (17) of 75 genes, 3.4 (SD 1.6) million reads are sequenced on average per sample. 55% (SD 10%) of these map to off-target regions of the genome. This gives a mean read depth in off-target regions of 0.065 (SD 0.044). In the exome samples that we used as a benchmarking data set there are an average of 76 (SD 20) million reads per sample with 20.3% (SD 6.6%) off-target, equating to mean read depth of 0.52 (SD 0.20) in off-target regions. This compares to a typical genome sequencing experiment where sufficient reads are sequenced to give >30X mean coverage across the genome.

### SavvyCNV can call off-target CNVs from targeted panels

To evaluate SavvyCNV’s ability to call off-target CNVs accurately from targeted panel data we benchmarked its performance against a truth set (see Methods) and compared it to five other tools for calling CNVs: GATK gCNV(18), DeCON(22), EXCAVATOR2(14), CNVkit(15), and CopywriteR(16). To prevent bias due to software configuration tuning, we ran all six tools with multiple configurations, and plotted the best results for each tool on a precision-recall graph (Figure 1). The best recall (sensitivity) where precision is at least 50% is shown in Table 1.

**Figure 1.**
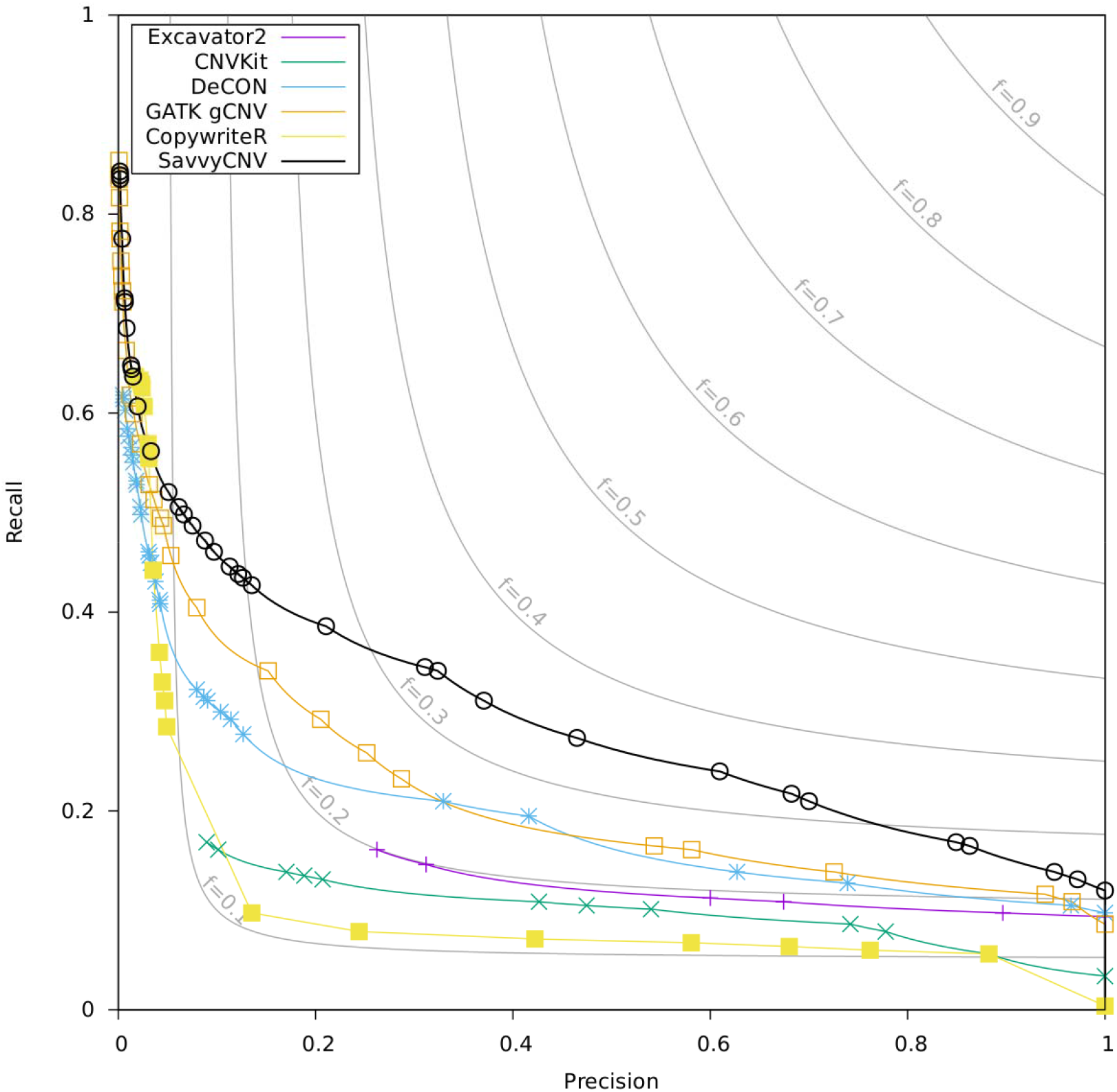
Benchmarking off-target CNV calling from targeted panel data. The data points on the plot are generated by a parameter sweep for each tool and show the precision and recall that can be achieved with each tool. The f statistic is the harmonic mean of precision and recall (see methods for details).

**Table 1.** Benchmarking off-target CNV calling from targeted panel data. The table shows the performance of the different CNV calling software based on the size of the CNV. The tools were run with multiple different parameters. For this comparison, we have selected the configuration for each tool that provides the highest recall with a precision of at least 50%. More variants may be detected by each tool with different configuration, but with precision less than 50%.

| Size | CNVs | Software | True positives | False positives | Recall | Precision |
| --- | --- | --- | --- | --- | --- | --- |
| All sizes | 267 | SavvyCNV | 68 | 67 | 25.5% | 50.4% |
|  |  | GATK gCNV | 40 | 31 | 14.9% | 56.3% |
|  |  | DeCON | 41 | 40 | 15.4% | 50.6% |
|  |  | Excavator2 | 30 | 20 | 11.2% | 60.0% |
|  |  | CnvKit | 27 | 23 | 10.1% | 54.0% |
|  |  | CopywriteR | 18 | 13 | 6.7% | 58.1% |
| >=1Mb<br>(including<br>>=5Mb) | 42 | SavvyCNV | 41 | 11 | 97.6% | 78.8% |
|  |  | GATK gCNV | 39 | 32 | 92.9% | 54.9% |
|  |  | DeCON | 36 | 31 | 85.7% | 53.7% |
|  |  | Excavator2 | 29 | 19 | 69.0% | 60.4% |
|  |  | CnvKit | 26 | 24 | 61.9% | 52.0% |
|  |  | CopywriteR | 18 | 13 | 42.9% | 58.1% |
| >=5Mb | 26 | SavvyCNV | 26 | 0 | 100% | 100% |
|  |  | GATK gCNV | 26 | 3 | 100% | 89.7% |
|  |  | DeCON | 26 | 19 | 100% | 74.3% |
|  |  | Excavator2 | 26 | 3 | 100% | 89.7% |
|  |  | CnvKit | 25 | 23 | 96.2% | 52.1% |
|  |  | CopywriteR | 17 | 8 | 65.4% | 68.0% |

All tools except CopywriteR called all of the CNVs larger than 5Mb (although not necessarily with precision of at least 50%), however only SavvyCNV did so without any false positive calls. All CNVs larger than 1Mb were called by SavvyCNV, GATK gCNV, and DeCON (with precision less than 50%), although SavvyCNV called the most (97.6%) at a precision of at least 50% (as in table 1). For all CNVs, SavvyCNV had the highest recall (25.5%) with precision of at least 50%. For all three CNV size categories, SavvyCNV had the greatest detection power. It can call CNVs that are larger than 1Mb from off-target reads from a targeted panel with good recall (97.6%) and precision (78.8%).

### SavvyCNV can call on-target CNVs from targeted panels

To evaluate the performance of SavvyCNV at calling CNVs from on-target data we used the ICR96 validation series(21) and compared its performance to GATK gCNV, DeCON, and CNVkit. ICR96 is a set of 96 samples sequenced using a small targeted sequencing panel (TruSight Cancer Panel v2, 100 genes), with exon CNVs detected independently using MLPA (25 single-exon CNVs, 43 multi-exon CNVs, and 1752 normal copy number genes). SavvyCNV had the highest recall for precision of at least 50% though GATK gCNV and DeCON also performed well - these 3 tools had a recall >95% (Table 2). Precision can only be compared between tools if recall is identical. While GATK gCNV achieves 85.7% precision at its highest recall of 97.1%, SavvyCNV has a precision of 93.0% at the same recall (this is shown in Figure 2). DeCON was the next-best performing tool while CnvKit did not call the majority of CNVs. Excavator2 did not run on this data set. Figure 2 shows the recall and precision of the four tools. SavvyCNV was the only tool capable of detecting all the CNVs although only with a precision of 29.1%.

**Table 2.** Benchmarking on-target CNV calling from the ICR96 targeted panel data. The table shows the performance of the different CNV calling software based on the size of the CNV. The tools were run with multiple different parameters. For this comparison, we have selected the configuration for each tool that provides the highest recall with a precision of at least 50%.

| Size | CNVs | Software | True positives | False positives | Recall | Precision |
| --- | --- | --- | --- | --- | --- | --- |
| All (single exon and multi-exon) | 68 | SavvyCNV | 67 | 42 | 98.5% | 61.5% |
|  |  | GATK gCNV | 66 | 11 | 97.1% | 85.7% |
|  |  | DeCON | 65 | 65 | 95.6% | 50.0% |
|  |  | CnvKit | 12 | 12 | 17.6% | 50.0% |
| Multi-exon | 43 | SavvyCNV | 43 | 11 | 100% | 79.6% |
|  |  | GATK gCNV | 43 | 11 | 100% | 79.6% |
|  |  | DeCON | 42 | 14 | 97.7% | 75.0% |
|  |  | CnvKit | 10 | 8 | 23.2% | 55.5% |

Two of the CNVs within the ICR96 dataset cover less than a complete exon and have one breakpoint within the targeted region. These two CNVs are the hardest to detect by read-depth methods, as the read depth is only altered over a fraction of the exon area. Both CNVs are detected only by SavvyCNV, even when the highest sensitivity settings are used with the other CNV callers.

Multi-exon CNVs are easier to detect than single-exon CNVs. SavvyCNV, GATK gCNV, and DeCON can detect all 43 multi-exon CNVs, although only SavvyCNV and GATK gCNV did this with a precision of at least 50%.

**Figure 2.**
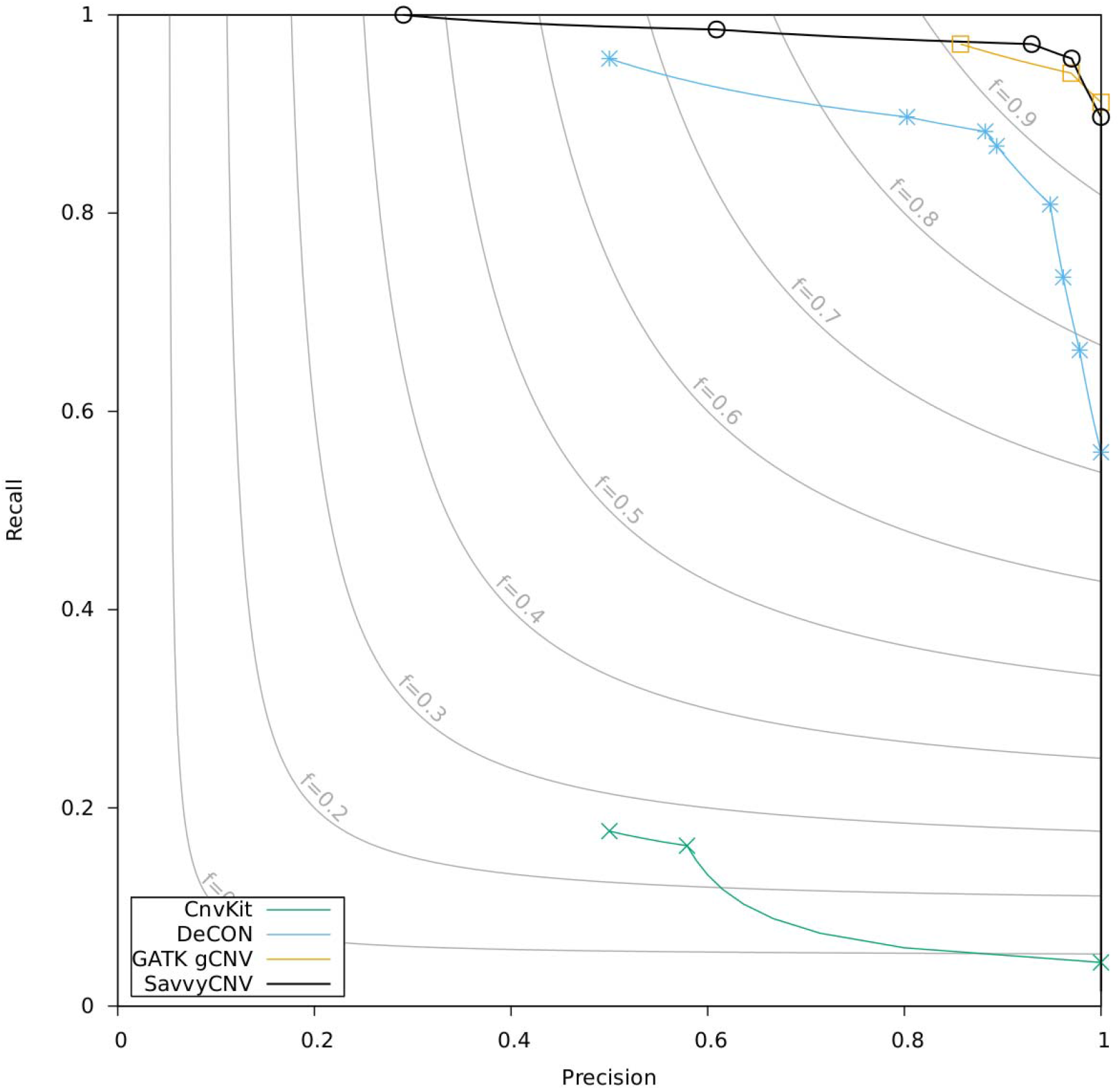
Benchmarking on-target CNV calling from targeted panel data. The data points on the plot are generated by a parameter sweep for each tool and show the precision and recall that can be achieved with each tool. The f statistic is the harmonic mean of precision and recall (see methods for details).

### SavvyCNV can call off-target CNVs from exome data

To assess SavvyCNV’s ability to call CNVs from off-target reads generated by exome sequencing we benchmarked it against a truth set (see Methods) and compared its performance to GATK gCNV, DeCON, EXCAVATOR2, CNVkit, and CopywriteR. The best recall where precision is at least 50% is shown in Table 3 for two different size categories, and recall/precision is shown in Figure 3 for all CNVs.

SavvyCNV was the best performing tool on this data set, able to call 86.7% of the CNVs with at least 50% precision, while the next best tool (DeCON) called 46.7% of CNVs with at least 50% precision. The chief difference between the performances of the tools is SavvyCNV’s ability to call CNVs smaller than 200kb. SavvyCNV is able to call an additional 30 CNVs that are smaller than 200kB at >=50% precision while GATK gCNV, EXCAVATOR2, and CNVkit call no true CNVs smaller than 200kB, DeCON calls 10, and CopywriteR calls 4.

**Figure 3.**
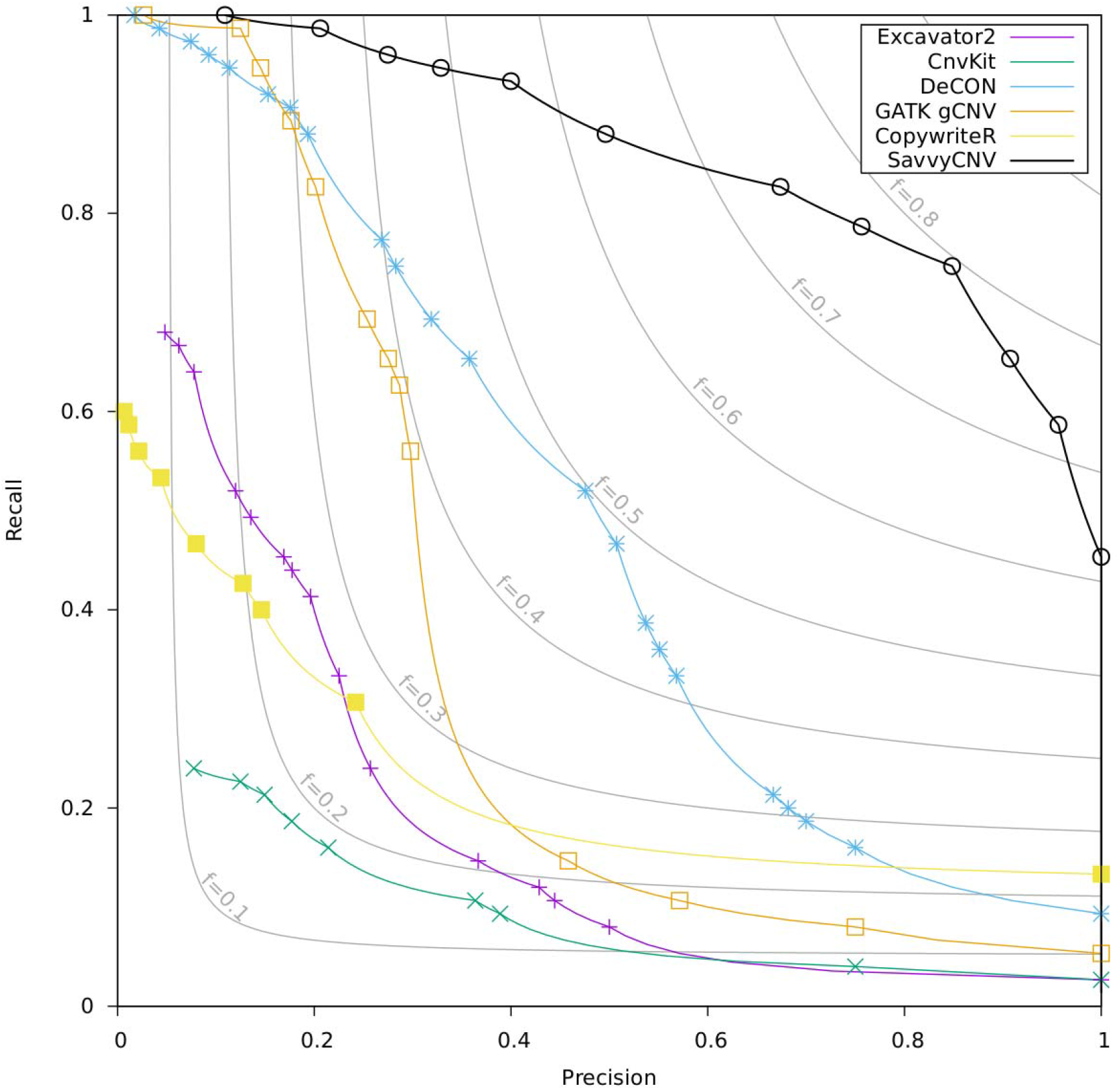
Benchmarking off-target CNV calling from exome data. The data points on the plot are generated by a parameter sweep for each tool and show the precision and recall that can be achieved with each tool. The f statistic is the harmonic mean of precision and recall (see methods for details).

**Table 3.** Benchmarking on-target CNV calling from the exome data. The table shows the performance of the different CNV calling software based on the size of the CNV. The tools were run with multiple different parameters. For this comparison, we have selected the configuration for each tool that provides the highest recall with a precision of at least 50%.

| Size | CNVs | Software | True positives | False positives | Recall | Precision |
| --- | --- | --- | --- | --- | --- | --- |
| All sizes | 75 | SavvyCNV | 65 | 58 | 86.7% | 52.8% |
|  |  | GATK gCNV | 9 | 9 | 12.0% | 50.0% |
|  |  | DeCON | 35 | 34 | 46.7% | 50.7% |
|  |  | Excavator2 | 6 | 6 | 8.0% | 50.0% |
|  |  | CnvKit | 3 | 1 | 4.0% | 75.0% |
|  |  | CopywriteR | 10 | 0 | 13.3% | 100% |
| >=200kb | 40 | SavvyCNV | 35 | 34 | 87.5% | 50.7% |
|  |  | GATK gCNV | 9 | 9 | 22.5% | 50.0% |
|  |  | DeCON | 25 | 24 | 62.5% | 51.0% |
|  |  | Excavator2 | 6 | 6 | 15.0% | 50.0% |
|  |  | CnvKit | 3 | 1 | 7.5% | 75.0% |
|  |  | CopywriteR | 6 | 0 | 15.0% | 100% |

### SavvyCNV can detect clinically relevant CNVs

Having validated the ability of SavvyCNV to call CNVs from off-target reads we proceeded to screen for CNVs in our cohort of targeted panel samples from patients referred for genetic testing to identify the cause of their diabetes or hyperinsulinism(17). We were able to detect 11 clinically relevant CNVs both within and outside of the targeted regions (Table 4). Of these, 4 provided a new genetic diagnosis for diabetes/hyperinsulinism (rows 1-4 in table 4), providing information which will guide clinical management and allow accurate counselling on recurrence risk in family members and future offspring. The remaining 7 CNVs (rows 5-11 in table 4) confirmed clinically-reported diagnoses unrelated to the diabetes/hyperinsulinism. These findings demonstrate the ability of SavvyCNV to detect clinically relevant CNVs and aneuploidies from off-target data from a small targeted panel.

**Table 4.** – Clinically-relevant CNVs detected.

| Row | CNV detected | CNV size | Clinical confirmation | Reason for referral | Clinical implications |
| --- | --- | --- | --- | --- | --- |
| 1 | Chr8:6,800,000-11,800,000 deletion | 5Mb | Deletion includes *GATA4*, causative of the patient's neonatal diabetes and additional features. | Diabetes | Genetic diagnosis of monogenic diabetes (23). |
| 2 | Chr8:8,000,000-10,400,000 duplication<br>8:10,600,000-12,000,000 deletion | 2.4Mb and 1.4Mb | Deletion includes *GATA4*, causative of the patient's neonatal diabetes and additional features. | Diabetes | Genetic diagnosis of monogenic diabetes (23). |
| 3 | Chr18:19,400,000-21,800,000 deletion | 2.4Mb | Deletion includes *GATA6*, causative of the patient's neonatal diabetes and additional features. | Diabetes | Genetic diagnosis of monogenic diabetes (24). |
| 4 | ChrX:0-57,000,000 deletion<br>X:76,200,000-155,400,000 deletion | 57Mb and 79.2Mb | Ring X chromosome confirmed by cytogenetics. | Hyperinsulinism | Genetic diagnosis of Turner syndrome, a known cause of hyperinsulinism (25). |
| 5 | Chr21:9,400,000-48,200,000 duplication | Chromosome | Patient known to have Down Syndrome at referral. | Hyperinsulinism | Confirms the diagnosis of Down syndrome. |
| 6 | Chr21:10,000,000-48,200,000 duplication | Chromosome | Patient known to have Down Syndrome at referral. | Diabetes | Confirms the diagnosis of Down syndrome (26). |
| 7 | Chr21:11,000,000-48,200,000 duplication | Chromosome | Patient known to have Down Syndrome at referral. | Hyperinsulinism | Confirms the diagnosis of Down syndrome. |
| 8 | Chr21:14,400,000-48,200,000 duplication | Chromosome | Patient known to have Down Syndrome at referral. | Hyperinsulinism | Confirms the diagnosis of Down syndrome. |
| 9 | Chr22:18,800,000-21,600,000 deletion | 1.8Mb | Confirmed by array CGH. | Diabetes | Provided the diagnosis of DiGeorge syndrome. |
| 10 | Chr22:18,800,000-21,600,000 deletion | 1.8Mb | Patient known to have DiGeorge syndrome at referral. | Hyperinsulinism | Confirms the diagnosis of DiGeorge syndrome. |
| 11 | ChrX:0-155,400,000 duplication | Chromosome | Patient known to have XXX syndrome at referral. | Diabetes | Confirms the diagnosis of XXX syndrome. |

## Discussion

### SavvyCNV can detect CNVs genome-wide from off-target reads

We benchmarked SavvyCNV on its ability to call off-target and on-target CNVs from targeted panel and exome sequencing data. This new tool outperformed five existing tools, three of which (CNVkit(15), Excavator2 (14), and CopywriteR(16)) were specifically designed to call off-target CNVs. GATK gCNV performed similarly to SavvyCNV in the on-target (ICR96) analysis. However, SavvyCNV considerably outperforms all other tools in the off-target analyses.

SavvyCNV finds the greatest number of true positive CNVs in all data sets while other tools did not call certain CNVs. For example, the two partial exon CNVs in the on-target (ICR96) data set are detected only by SavvyCNV. This is likely because of the improved error correction and error modelling that is incorporated into SavvyCNV over existing tools. SavvyCNV uses singular vector decomposition to reduce noise. CNVkit, EXCAVATOR2 and CopywriteR only correct for GC content, while GATK gCNV uses Bayesian principle component analysis (https://www.broadinstitute.org/videos/scalable-bayesian-model-copy-number-variation-bayesian-pca), and DeCON uses sample matching (it searches for samples in the control set that have a similar noise profile). Unlike the other tools tested, CopywriteR does not normalise against other samples but excludes on-target reads to make read counts representative of the true copy number. Supplementary figure 1 demonstrates how error correction improves the recall and precision. Supplementary figure 2 shows that the bespoke error model used by SavvyCNV (see supplementary methods) performed better than the Poisson error model used by the other tools, as this uses the error information available from having multiple control samples.

SavvyCNV had a higher precision than other tools when calling off-target CNVs, an important consideration in diagnostic and research laboratories as if false positives are reduced, fewer CNVs will require orthogonal testing to identify the true positive results. Many of the false positives produced by DeCON, CNVKit, and EXCAVATOR2 have a read depth ratio indicating that they are either mosaic CNVs or random noise. The prior probability is overwhelmingly that these are random noise. This is why the default for SavvyCNV and GATK gCNV is to call only non-mosaic CNVs as this hugely reduces the number of false positives called. Mosaic CNV calling can be enabled in SavvyCNV for projects where it is applicable. Supplementary figure 3 demonstrates the improvement in precision of the default mode compared to the mode that includes calling of mosaic CNVs.

### Estimates of precision and recall rely on the quality of the truth set

On-target CNV calling from a targeted panel was tested on the ICR96 data set in which the truth set was verified by MLPA. The truth sets for the off-target CNV calling from targeted panel and exome sequence data were generated from CNV calls from genome sequencing data. Genome sequencing has a much higher coverage than that generated from only off-target reads which allows CNVs to be called more accurately enabling them to be used as a truth set. GenomeStrip(20) was used to call the truth set as it was designed to call CNVs from genome data and was not one of the tools under examination in this study. However, it is possible that there could be some false positive and negative calls in the truth set. This would lower the precision and recall of the tools under examination but should not bias the results in favour of a particular tool.

### Sensitivity depends on the size of the CNV

Smaller CNVs are harder for all software to detect. For all tools tested the larger the CNV the better the precision and recall, however SavvyCNV performs better than the other tools tested. SavvyCNV detects CNVs above 1Mb with 100% recall in off-target data from both targeted panel and exome data.

### CNV calling can be optimised for precision or recall by adjusting configuration

When calling CNVs, precision and recall are a trade-off; high recall will maximise the number of true CNVs that are called, with the consequence that it also reduces precision resulting in a large number of false positive CNV calls. Different precision levels are appropriate in different situations, influenced both by the experimental methodology and the aims of the project. When calling CNVs on-target on a small gene panel there will be fewer false positive calls generated due to the smaller target area thus it may be preferable to adjust settings to enable a higher recall at the cost of a lower precision. This could also be true in a clinical context where the most important aim is to not miss a true causative variant. In contrast, when calling CNVs genome-wide in a gene-agnostic approach such as genome sequencing, a higher precision is likely to be desirable to avoid generating an unmanageably long list of CNVs. The user can choose their preferred settings for SavvyCNV for different project requirements.

### Off-target CNV calling is ‘free’ data and increases diagnostic yield

SavvyCNV utilises data already generated by targeted panel and exome tests. These tests are carried out in order to detect single nucleotide variants and small insertions or deletions (<50 base pairs). In some laboratories CNVs are also detected within the targeted regions using CNV calling software while other laboratories use array-CGH or MLPA to detect CNVs. Using SavvyCNV allows CNVs to be detected not just within the targeted regions but allows genome-wide CNV calling. This will provide a genetic diagnosis for more patients, increasing the diagnostic yield of these tests. We have demonstrated the ability to find relevant genetic diagnoses using off-target CNV calling from our small targeted panel. Existing data can be reanalysed with our method to reveal additional CNVs. As an illustration of this, two of the CNVs in the ICR96 data set were found to actually be large CNVs (15Mb and 56Mb), which may have clinical implications beyond the targeted gene.

SavvyCNV calls CNVs from off-target reads from exomes and small targeted gene panels with high precision and recall, and performs better than existing tools including those designed for off-target CNV calling. Calling CNVs from off-target reads is exploiting ‘free’ data to increase the diagnostic yield of targeted panel and exome sequencing tests and reveal important biological findings.

## Data Availability

SavvyCNV is available for download from https://github.com/rdemolgen/SavvySuite The targeted panel and exome sequencing data analysed during the current study is not publicly available due to patient confidentially.

## Funding

SE is the recipient of a Wellcome Senior Investigator award (grant number WT098395/Z/12/Z). SEF has a Sir Henry Dale Fellowship jointly funded by the Wellcome Trust and the Royal Society (grant number: 105636/Z/14/Z). KAP holds a post-doctoral fellowship from the Wellcome trust (110082/Z/15/Z). This work was undertaken as part of the Transforming Genetic Medicine Initiative funded by the Wellcome Trust.

The funding bodies had no role in the design of the study or the collection, analysis or interpretation of data.

## Conflict of interest

The authors declare that they have no conflicts of interests

## Acknowledgements

The authors would like to thank Professor Andrew Hattersley, Dr Jayne Houghton and Andrew Parish (Department of Molecular Genetics, Royal Devon & Exeter NHS Foundation Trust, Exeter, UK) for their technical assistance and advice. The authors would like to acknowledge the use of the University of Exeter High-Performance Computing (HPC) facility in carrying out this work.

## Notes

#### Summary of Updates

An analysis of the performance of CopywriteR on the three data set has been added. Many thanks for the comments bringing this paper/tool to our attention.

