## Supplementary information for "SavvyCNV: genome-wide CNV calling from off-target reads"

### Supplementary figures:

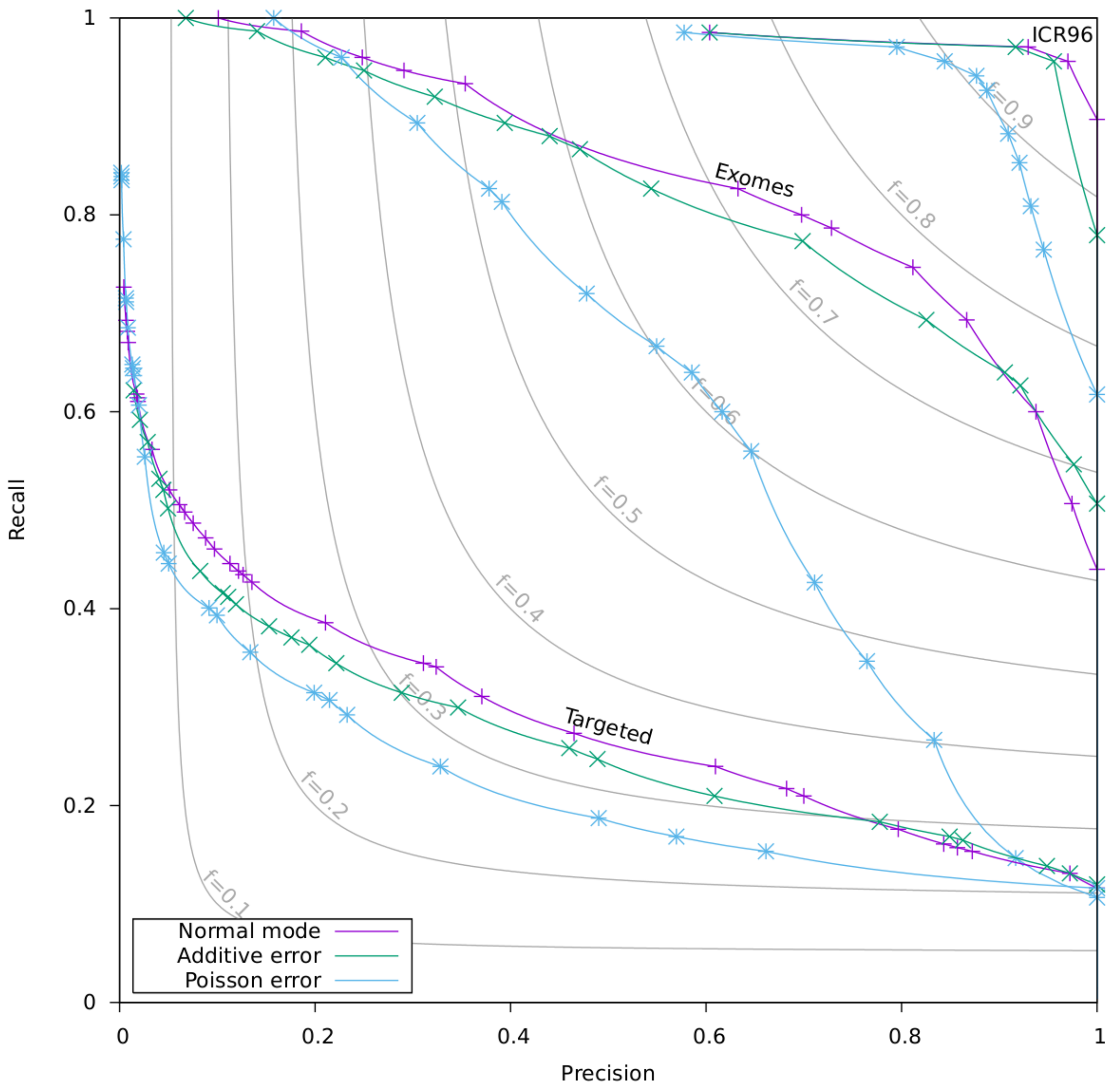


Supplementary Figure 1: Shows the improvement in precision/recall due to the error modelling strategy used by SavvyCNV. By default, SavvyCNV estimates the error in each genomic location in each sample by combining the average error in each sample with the average error in each genomic location. The default configuration does this by multiplying the two error values. An alternative is to add the two error values. Other software assumes that the error is Poisson in nature, and can therefore be calculated from the read depth – the results using this assumption are also plotted. The error estimate is used to determine whether the read depth in a genomic location is significantly outside the range expected for normal copy number. In reality, the error is greater than the Poisson estimate in some genomic locations, which makes error modelling beneficial.

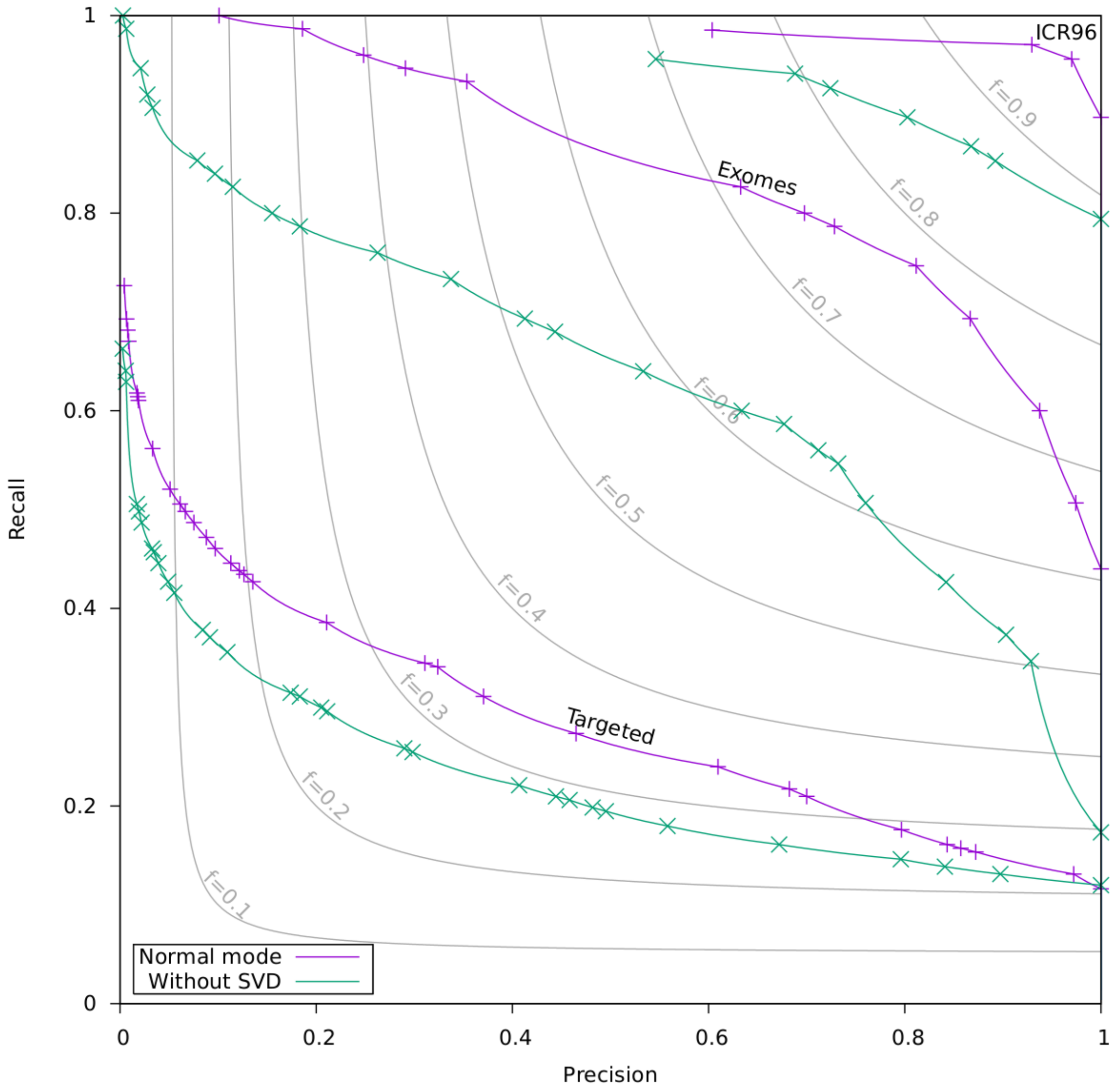


Supplementary Figure 2: Shows the improvement in precision/recall due to the error correction strategy used by SavvyCNV, for all three data sets. By default, SavvyCNV uses singular vector decomposition (SVD), which identifies biases common to multiple samples, which can be caused by differences in sample handling or chemistry. The "normal" line shows the default configuration, while the "No SVD" line is with SVD switched off.

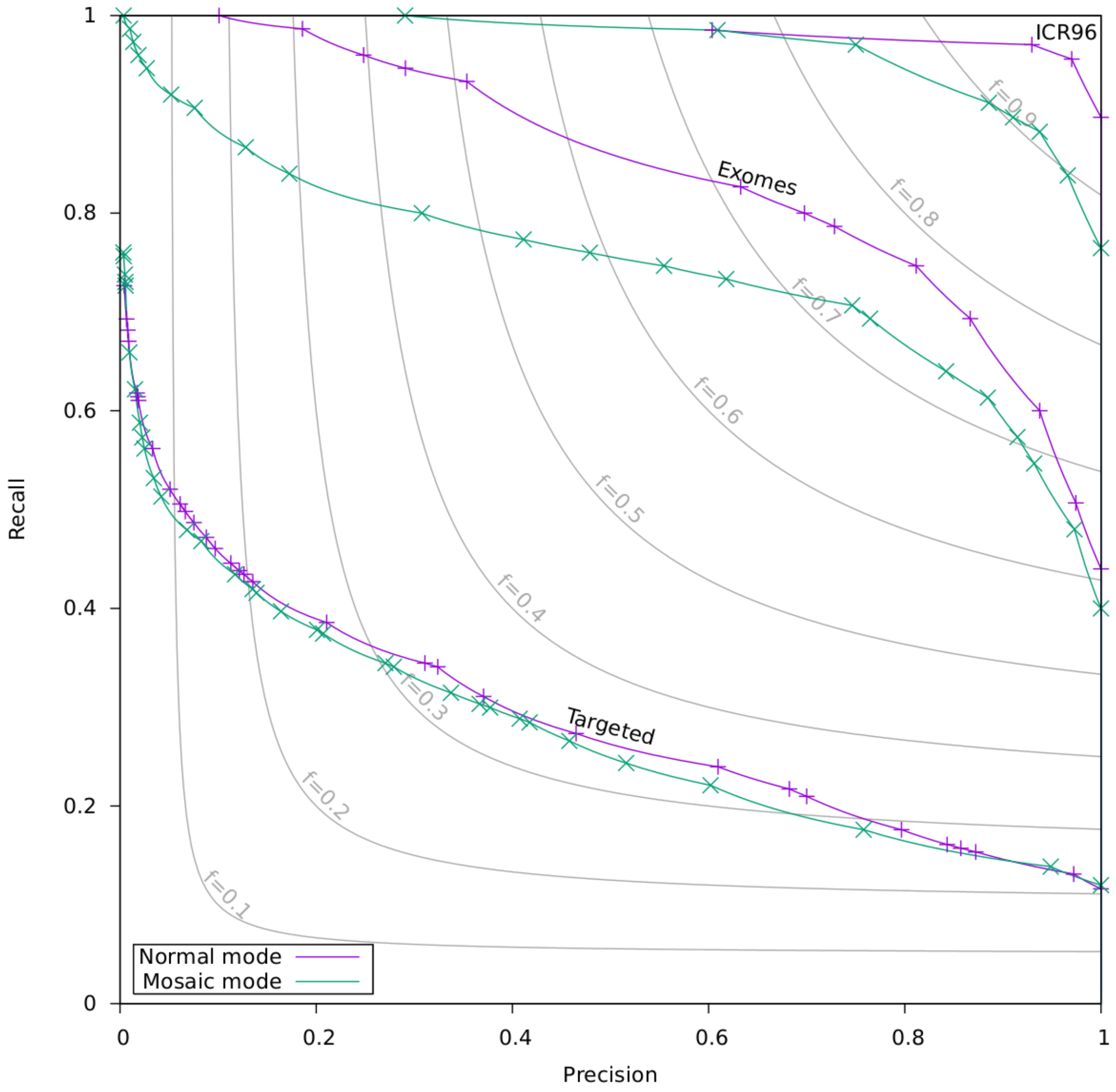


Supplementary Figure 3: Shows the improvement in precision/recall due to the default integer copy number strategy used by SavvyCNV. By default, SavvyCNV assumes that CNVs are not mosaic. This allows it to reduce false positives by requiring more evidence in the normalised read depth values. In mosaic mode (and DeCON, Excavator2, and CNVKit), a normalised read depth must differ from 1.0 by more than the error to be evidence of a CNV. In SavvyCNV's default mode (and GATK gCNV), the read depth must also be closer to 0.5 or 1.5 (representing a whole heterozygous deletion or duplication) than 1.0 to be evidence of a CNV.
